# Dual medial prefrontal cortex and hippocampus projecting neurons in the paraventricular nucleus of the thalamus

**DOI:** 10.1101/2021.10.02.462854

**Authors:** Tatiana D. Viena, Gabriela E. Rasch, Timothy A. Allen

**Author notes:** Corresponding Author: Timothy A. Allen, PhD, Department of Psychology, Florida International University, 11200 SW 8^th^ Street, Miami, FL, 33199, Website: http://allenlab.fiu.edu/.

## Abstract

The paraventricular nucleus (PVT) of the midline thalamus is a critical higher-order cortico-thalamo-cortical integration site that plays a critical role in various behaviors including reward seeking, cue saliency, and emotional memory. Anatomical studies have shown that PVT projects to both medial prefrontal cortex (mPFC) and hippocampus (HC). However, dual mPFC-HC projecting neurons which could serve a role in synchronizing mPFC and HC activity during PVT-dependent behaviors, have not been explored. Here we used a dual retrograde adenoassociated virus (AAV) tracing approach to characterize the location and proportion of different projection populations that send collaterals to mPFC and/or ventral hippocampus (vHC). Additionally, we examined the distribution of calcium binding proteins calretinin (CR) and calbindin (CB) with respect to these projection populations PVT. We found that PVT contains separate populations of cells that project to mPFC, vHC, and those that innervate both regions. Interestingly, dual mPFC-HC projecting cells expressed neither CR or CB. Topographically, mPFC- and vHC-projecting CB^+^ and CR^+^ cells clustered around dual projecting neurons in PVT. These results are consistent with the features of dual mPFC-vHC projecting cells in the nucleus reuniens (RE) and suggestive of a functional mPFC-PVT-vHC system that may support mPFC-vHC interactions in PVT-dependent behaviors.

## Introduction

Interactions between the medial prefrontal cortex (mPFC) and the hippocampus (HC) support various cognitive functions at the intersection of learning, memory, emotion, and executive function (Jones and Wilson 2005; Churchwell and Kesner 2011; Ito et al. 2015; Eichenbaum 2017). For example, the connections between the mPFC and HC are critical to flexible spatial memory (Wirt and Hyman 2017; Avigan et al. 2020), but this is not always found with simpler forms of spatial memory (Aggleton and Brown 1999; Yoon et al. 2008).

The midline thalamus bi-directionally connects mPFC and HC forming higher order cortico-thalamo-cortical circuits supporting behaviors that depend on mPFC, HC and mPFC-HC interactions (Huang et al. 2006; Vertes et al. 2007, 2015; Kirouac 2015; Choi and McNally 2017; Dolleman-van der Weel et al. 2019). Anatomically, the midline thalamus is divided into four principal nuclei including the dorsal paraventricular (PVT) and paratenial (PT) nuclei and the ventral rhomboid (RH) and reuniens (RE) nuclei. These are generally thought of as composing separate thalamocortical circuits within the limbic system (Groenewegen and Berendse 1994; Van der Werf et al. 2002; Vertes et al. 2015; Kirouac 2015). RE is most often implicated in HC-dependent learning and memory (Cholvin et al. 2013; Cassel et al. 2013, 2021; Viena et al. 2018; Jayachandran et al. 2019; Dolleman-van der Weel et al. 2019), whereas PVT is most often implicated in amygdala-dependent emotional memory (Padilla-Coreano et al. 2012; Do-Monte et al. 2015; Penzo et al. 2015) and nucleus accumbens related addictive behaviors (Martin-Fardon and Boutrel 2012; Zhu et al. 2016; Neumann et al. 2016).

Recent papers have emphasized the importance of thalamic cells that project to both mPFC and HC which have exclusively focused on RE (Hoover and Vertes 2012; Varela et al. 2014; Viena et al. 2021). These dual projecting cells are thought to be important in forming a unique mPFC←RE→vHC circuit providing synchronizing activity thus facilitating mPFC-HC interactions. Thus, dual projecting neurons are enabling of permissive network wide activity modes suitable for the acquisition, flexible retrieval, and flexible use of memory in support of adaptive behavior (Hallock et al. 2016; Zimmerman and Grace 2016; Eichenbaum 2017; Ferraris et al. 2018; Dolleman-van der Weel et al. 2019; Hauer et al. 2019; Schultheiss et al. 2020; Viena et al. 2021).

Here we focused on PVT given it also has anatomical connections with mPFC and HC (Su and Bentivoglio 1990; Vertes and Hoover 2008; Gergues et al. 2020). PVT spans the entire rostrocaudal extent of the midline thalamus and is positioned just ventral to the dorsal third ventricle (3V), medial to the paratenial nucleus (PT) in anterior levels and to the mediodorsal nucleus (MD) at posterior levels (Groenewegen and Berendse 1994) within the thalamus. PVT exhibits dense projections to mPFC, ventral CA1 (in slm), subiculum, nucleus accumbens and central/basal nuclei of the amygdala (Su and Bentivoglio 1990; Vertes and Hoover 2008). Recently, two anatomically and functionally distinct cell populations within rodent PVT that project to mPFC were found to represent independent circuits biased towards the anterior or posterior areas of PVT, as determined by gene expression (Gao et al. 2020).

Behaviorally, PVT projections have been extensively studied in the context of reward seeking, addictive behaviors, such as context-induced addiction and relapse memories, cue-saliency, and emotional processing (James et al. 2010; Heydendael et al. 2011; Penzo et al. 2015; Kirouac 2015; Neumann et al. 2016; Hill-Bowen et al. 2020). In emotional memory, PVT projections to the prelimbic (PL) and infralimbic (IL) cortices of mPFC are important for the retrieval of fear memories (Huang et al. 2006; Padilla-Coreano et al. 2012; Do-Monte et al. 2015). PVT also projects to the hippocampus, specifically to ventral CA1 (Su and Bentivoglio, 1990; Gergues et al., 2020) and ventral subiculum (Moga et al. 1995; Otake and Nakamura 1998; Vertes and Hoover 2008; Li and Kirouac 2012), and HC→PVT projections are implicated in associative learning (Haight and Flagel 2014; Dong et al. 2015; Choi and McNally 2017; Zhu et al. 2018). The vHC, known for its role in anxiety and emotionally charged contextual memories, (Moser and Moser 1998; Bannerman et al. 2003; Squire et al. 2004; Xu et al. 2016), involves an newly identified an anatomical circuit from PVT→vCA1→mPFC (Gergues et al. 2020).

Despite what seems to be a clear involvement of PVT in memory processes, the neural circuits associated with PVT remain understudied. Given known PVT projections to mPFC and HC, individual PVT neurons may send projections to both mPFC and HC, similar to RE, or these may be entirely segregated projection populations. Here we used a dual retrograde adenoassociated virus (AAV) approach to look at PVT neurons that project to mPFC and HC, as done previously in RE (Hoover and Vertes 2012; Varela et al. 2014; Viena et al. 2021). We delivered bilateral retrograde AAV into ventral CA1 (vCA1; retro AAV-GFP) and to prelimbic/infralimbic (PL/IL; retro AAV-tdTomato) regions of the rodent mPFC. We also counterstained for the calcium binding proteins calretinin (CR) and calbindin (CB), given that both the dorsal and ventral midline thalamus are similarly organized into cell density zones that preferentially express one or both calcium binding proteins (Viena et al. 2021). Specifically, we sought to identify neuronal populations within PVT that 1) independently project to vHC and mPFC, or 2) exhibit collaterals to both mPFC and vHC and 3) describe their topography in relation to CR and CB immuno-reactive positivity. The results showed PVT contains three projection populations with respect to the mPFC-HC system that project to mPFC, vHC or both mPFC and vHC. Notably the dual-projecting cells were located throughout its entire rostro-caudal axis of PVT. Like RE, these dual projecting cells do not express either CR or CB but single projecting CR^+^ and CB^+^ cells cluster around them. This newly identified mPFC←PVT→HC circuit may provide a synchronizing mechanism permissible of mPFC-HC interactions in PVT-dependent behaviors.

## Methods

### Subjects

Four Long-Evans rats (2 females, 2 males; 250-350g) were used for this experiment. Upon arrival, rats were individually housed and allowed to acclimate for ∼1 week. Rats had free access to food and water. All procedures were conducted in accordance with Florida International University Animal Care and Use Committee (FIU IACUC).

### Delivery of retrograde AAV vector through stereotaxic surgery

Glass micropipettes were made prior to surgery using a laser-based micropipette puller (P-2000, Shutter Instruments). The micropipettes were used in conjunction with Nanoject III microinjector (3-000-2030g/x, Drummonds Scientific). In order to minimize brain damage in injected regions, a fine long taper (∼10mm) was used for the vHC injections and a shorter taper (∼5mm) was used for mPFC injections. Retrograde AAV vectors AAV-CAG-tdTomato (59462-AAVrg; Addgene, MA) and AAV-CAG-GFP (Addgene, MA; 37825-AAVrg) were delivered bilaterally in mPFC (AP +3.0 mm, ML +/-1.2mm, DV -5.0mm, 50nL; -4.4mm, 150 nL; and -3.8mm, 200 nL) and vHC (AP -5.6mm, ML+/-5.85mm, DV -7.2 mm,100 nL; 6.8 mm, 200 nL, and 6.2 mm, 200 nL), respectively (see Fig. 1A). The total AAV vector volume (mPFC 0.4μL and vHC 0.5 μL) was delivered at a rate of 1 nL/sec and allowed to diffuse for 5 min after each injection. Viral vector incubation times were between 6-8 weeks post-surgery.

**Figure 1.**
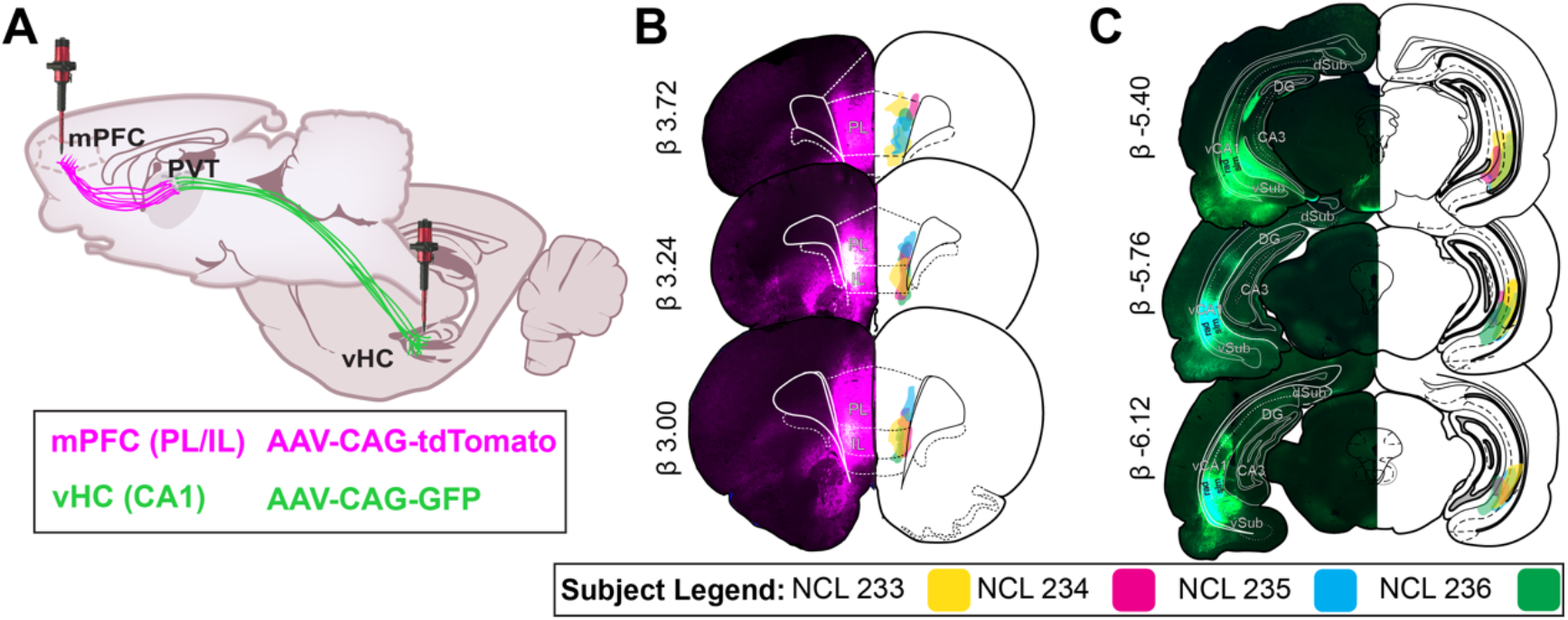
Retrograde AAV injections in mPFC and vHC. A) Paired bilateral injections of retrograde AAV-CAG-tdTomato (magenta) and retrograde AAV-CAG-GFP (green) delivered in mPFC and vHC for florescence protein expression of neurons in PVT. B) mPFC injection spread was confined to PL/IL regions, primarily layers V/VI (n = 4). Epifluorescence capture (4X) on the left depict the injection spread in PL/IL in one case (NCL 235). Retrograde labeling can be seen locally and in other frontal regions (VO, LO, ACC). On the right, a schematic representation of injection spread for each case across brain levels are illustrated. C) vHC injection spread was confined to CA1 molecular layers (n = 4). Epifluorescence capture (4X) on the left depict the injection spread in vCA1 in one case (NCL 235). Retrograde labeling can be seen locally in CA1, Sub and EC. On the right, a schematic representation of injection spread in vCA1 for each case across brain levels are illustrated. Abbreviations: AAV, adeno-associated virus; ACC, anterior cingulate cortex; CA1, CA1 subfield of hippocampus; CAG, chicken beta-Actin promoter; DG, dentate gryus; GFP, green fluorescent protein; HC, hippocampus; IL, infralimbic cortex; LO, lateral orbital cortex; mPFC, medial prefrontal cortex; PL, prelimbic cortex; rad, stratum radiatum; slm, stratum lacunosum moleculare; tdTomato, tandem dimer Tomato; vCA1, ventral CA1; VO, ventral orbital cortex.

### Histological tissue processing

Anesthetized rats were transcardially perfused with 200 mL heparinized phosphate-buffered saline (10 mL/min), followed by 250 mL of 4% paraformaldehyde (PFA, pH 7.4). The brains were removed, post-fixed for 24 hr in 4% PFA, and then transferred to a 30% sucrose solution until saturated. The frozen brains were sliced into coronal sections of 40μm using a cryostat (Leica CM 3050S). Brain tissue was preserved in 0.1 M phosphate buffer (PB) solution and protected from light with aluminum foil until mounted onto gelatin coated slides and cover slipped with Vectashield® antifade mounting medium with 4’,6-diamidino-2-phenylindole (DAPI) H-1200. Adjacent slices from another set of tissue underwent 3,3’-diaminobenzidine (DAB) reactions of calretinin (CR) or calbindin (CB) as described in Viena et al. (2021).

### Imaging brain slices

The effective spread of the viral particles was visualized using an epifluorescence microscope Olympus BX41 at 4x magnification. Atlas overlays from Swanson Rat Brain Atlas (2018) were used to define target regions using Adobe Illustrator ® (Adobe Systems Inc., San Jose, CA) for confirmation of injection sites and regional identification of PVT. Immunofluorescence from the retrogradely labeled PVT neurons was imaged using a confocal microscope (Olympus Fluoview FV-1200), at 10x magnification for an overview and 20x for quantifying virally infected neurons. The tdTomato viral vector was visualized using a red fluorescence cube; GFP containing viral vector was visualized using green fluorescence cube; and DAPI was visualized using blue fluorescence cube. After merging the red and green channels together, neurons that were yellow were classified as PVT neurons with axonal collateralizations to both mPFC and vHC. Mounted sections with DAB peroxidase were captured using an epifluorescence microscope Olympus BX41 at 20x magnification, in the same manner as the procedure described in Viena et al. (2021). Individual image captures were made in four channels: red (PVT→mPFC), green (PVT→vHC), blue (DAPI), and brightfield (DAB CR^+^ and DAB CB^+^ cells). Of note, the red channel was changed to magenta for visualization purposes (note that dual labeling shows white).

### Cell quantification

For each subject, cell counting was done across four PVT anterior-posterior levels: anterior (aPVT; AP -1.20mm), anteromedial (AP -1.72 mm), posteromedial (AP -2.52 mm), and posterior (pPVT; AP -3.00mm). We focused most of the analysis on the anterior and posterior levels as done by others (Li and Kirouac 2012; Do-Monte et al. 2015; Dong et al. 2017; Choi et al. 2019) but considered important to also characterize the transient medial regions of PVT as argued by Gao et al. (2020). When dual PVT mPFC-vHC projecting neurons were identified, we confirmed they were single neurons rather than overlapping neurons using z-stacks (0.5 um optical sections). Manual counts were performed with FIJI Image J and Adobe Photoshop ® (Adobe Systems Inc., San Jose, CA) in only one hemisphere to avoid intrinsic confounds and leveling differences. Cell counts were done manually by three trained experimenters and averaged. We validated the reliability of the cells counts by comparing the results between experimenters and their combined average. Cell counts from Experimenters 1, 2 and 3 were significantly and consistently correlated with the grand averaged count (Average v. Experimenter 1, r = .993; Average v. Experimenter 2, r = .994; Average v. Experimenter 3, r = .990; all p < .001). A test of inter-rater reliability was high among all experimenters’ counts and the grand average total (Cronbach’s α = .991). CellProfiler Software version 4.0.6 (cellprofiler.org) was used to perform automated DAPI counts using the validated and customized pipeline described in Viena et al. (2021).

### Calcium binding protein cell distances from dual mPFC-HC projecting cells

Adobe Photoshop was used with captures made on slides containing CR^+^ and CB^+^ DAB peroxidase-stained cells to transpose the brightfield image onto the merged fluorescent image. An area of 545 μm × 390 μm containing dual projecting cell clusters in the anterior and in posterior levels was selected for analysis. A separate layer marking the location of CR^+^ cells or CB^+^ cells with “+” symbols was created on FIJI ImageJ. Using a flattened image with the symbols, we assessed the co-expression between the virally infected cells and DAB-stained cells. Then, three trained experimenters measured the distance from the center of dual projecting cells to each of the DAB-stained cells surrounding them within a 100 μm radius circle (radial distance). All distance was measured using ImageJ’s ROI manager toolbox. Finally, we assessed the number of CR^+^ and CB^+^ DAB cells per every 10 μm radius area (up to 100 μm) from dual projecting cell locations to determine the density distribution (# of DAB cells/each radius area) of CR^+^ and CB^+^ cells with respect to the dual projecting cells.

### Data analysis

Data was tallied, graphed, and analyzed using Microsoft Excel, GraphPad Prism (version 9) and SPSS (version 27), respectively. We evaluated the percent cell density of the PVT→mPFC (magenta), PVT→vHC (green) and PVT dual projecting cells (white) across the four levels, where percent cell density was the ratio of counted cells over the number of DAPI nuclei (total number of cells in the region) at each level. One-way repeated measures analysis of variance (ANOVA) was used to determine if there were different mean percent cell densities across PVT cell populations. Following this, a linear regression was performed to determine the linear relationship between the frequency distance distribution of CR^+^ and CB^+^ cells with respect to dual projecting cells. Finally, we assessed radial cell area density distributions of CR^+^ and CB^+^ cells by running a comparison of fits between linear and exponential decay regressions, to model the type of distribution exhibited by these cells. An alpha of .05 was considered statistically significant for all analysis.

## Results

### Effective spread locations for retrograde AAV-tdTomato (mPFC) and retrograde AAV-GFP (vHC)

For all cases, we verified that the effective viral spread was localized within the PL/IL regions in mPFC (magenta) and ventral CA1 in vHC (green) as shown in Fig. 1B-C. For mPFC, fluorescence was confined to the deep layers (V/VI) of PL and IL regions. Local retrograde labeling is evident in frontal regions (VO, LO, ACC) (Fig. 1B), all of which are regions known to project to PL/IL. For vHC, fluorescence was mainly confined to CA1 molecular layers. Retrograde labeling can also be seen at that level locally in vCA1, as well as in entorhinal, ventral subiculum, and ventral dentate gyrus regions (Fig. 1C), which are regions known to contain neurons that project to vCA1.

### Dual mPFC-vHC projecting neurons found throughout the anterior-posterior extent of PVT

Evidence supporting the presence of dual mPFC-vHC projecting neurons was found throughout the anterior-posterior axis of PVT (Fig. 2Ai-Aiv). While posterior PVT cells are known to predominantly project to PL/IL (Vertes and Hoover 2008), at the anterior level of PVT (AP ∼ -1.20mm), we visualized both PVT mPFC-projecting, PVT vHC-projecting and PVT dual labeled neurons along the lateral borders of aPVT (Fig. 2 Ai). A higher density of mPFC-projecting cells was seen in dorsolateral regions at this level, but they also comingled with vHC-projecting cells along the lateral borders. This topography of mPFC- and vHC-projecting PVT cells is similar to RE and peri-reuniens relative to the location of the third ventricle. Dual projecting cells were observed along the entire lateral borders of PVT, typically surrounded by other mPFC- and vHC-projecting cells. In contrast, dual-projecting cells were sparse in midline regions where instead a dense mesh of fibers and puncta dominated the space. At this anterior level, a topographical separation of axonal processes and puncta was evident with mPFC-projecting cell axons and terminals being more abundant in dorsolateral portions, while vHC-projecting axonal processes and puncta was much more abundant in midline regions. This distribution further indicates that mPFC-projecting and vHC-projecting cells have a clear topographical organization within PVT (Fig. 2Ai). At this level, we focused on a group of mPFC-projecting, vHC-projecting and dual-projecting neurons at 100x magnification as shown on Fig. 2B. Here, it can be observed that mPFC- and vHC-projecting cells clustered around dual-projecting cells. Dense fibers and puncta from dual projecting cells, and stained nuclei by DAPI from other PVT cells (other than mPFC or vHC-projecting), were also observed in the same region (Fig. 2B).

**Figure 2.**
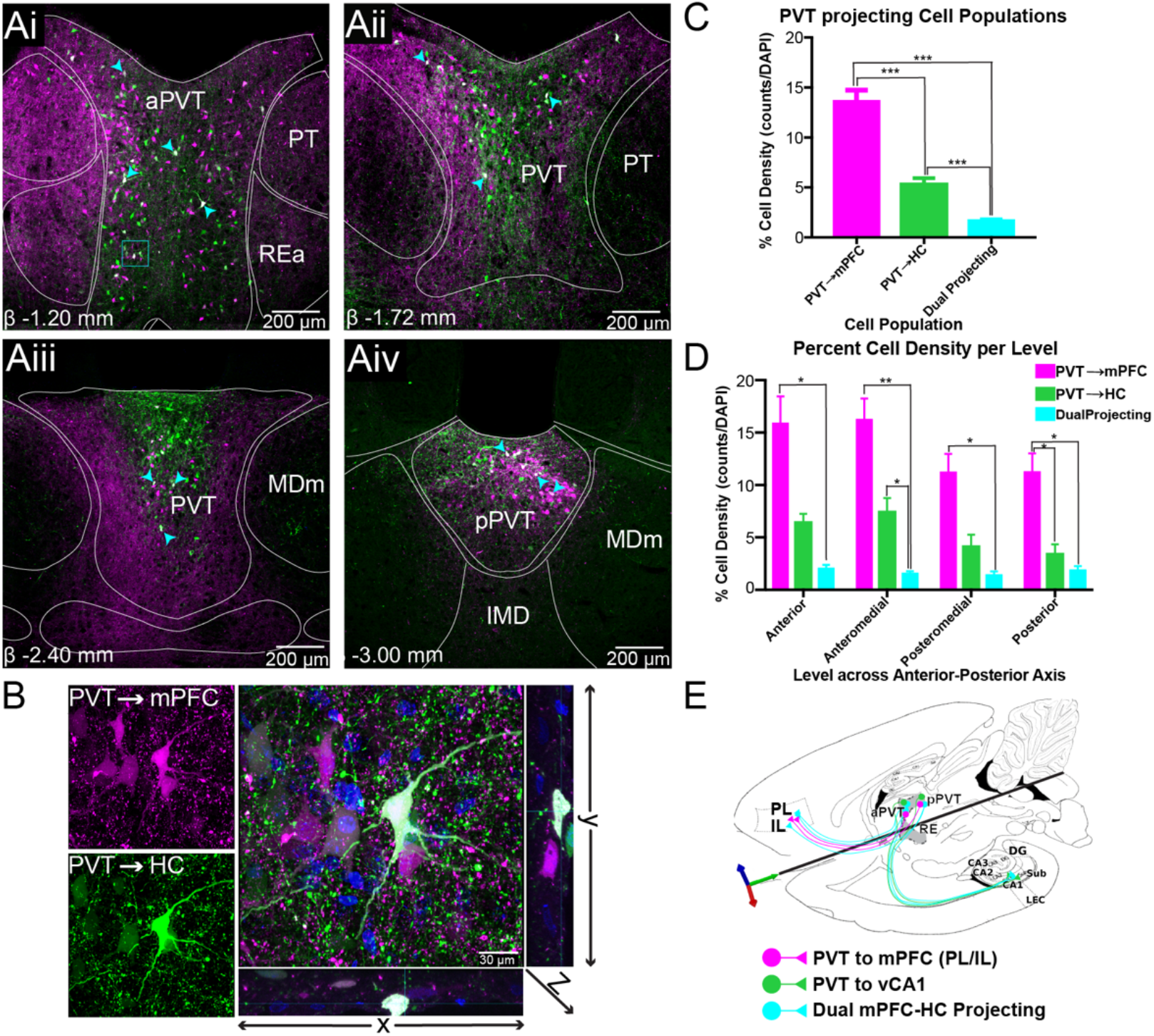
Dual mPFC-HC projecting cells are present throughout the anterior-posterior axis of PVT. A) Retrogradely labeled neurons from mPFC and HC in anterior (Ai), anteromedial (Aii), posteromedial (Aiii), and posterior levels (Aiv) of PVT. Images shown are not all from the same case. B) Confocal image (100x) of dual mPFC-HC projecting neuron in rostral PVT was obtained by merging the magenta (mPFC) and green (HC) channels, where dual labeled PVT cells are shown in white. An orthogonal view passing through the z-axis are shown to right (yz direction) and bottom (xz direction). C) mPFC-HC dual projecting percent cell density (# of cells/DAPI) summary in PVT (all levels) examined. PVT→mPFC cell density is larger than PVT→HC population. Dual cells represent 2% of all cells (DAPI) in PVT. All counts were performed in one hemisphere only. D) Percent cell density across each level examined. PVT→mPFC and PVT→HC cell density is higher in anterior levels but decreases in posterior levels. The dual mPFC-HC projecting cell density is consistent throughout all levels. E) A circuit conceptualization representing PVT cell projecting populations in the mPFC-HC circuit. These populations are present through the anterior-posterior levels of PVT. Image shown here is a transverse section containing mPFC, hippocampal formation, and midline thalamus. Abbreviations: DAPI, 4’,6-Diamidino-2-Phenylindole, Dihydrochloride; HC, hippocampus; mPFC, medial prefrontal cortex; PVT, paraventricular nucleus of the thalamus.

At anteromedial PVT levels (AP∼ -1.72mm), the cells redistributed topographically and had a “Y” shape that occupied mostly the dorsolateral and dorsomedial portions of the nucleus (Fig. 2Aii). These cells were surrounded by dense fibers and puncta similarly to that described in anterior levels. In the most ventral portion, there were little to no mPFC- or vHC-projecting cells. Dual-projecting cells were predominant in dorsolateral and dorsomedial regions spread amongst the band that overlaps with clusters of mPFC-projecting and vHC-projecting cells. At this level, we also observed many dual-projecting axons at the dorsolateral borders of PVT. PVT vHC-projecting axons were abundant in dorsomedial regions, while mPFC-projecting axons defined the lateral boundaries of the nucleus (Fig. 2Aii).

In posteromedial PVT levels (AP∼ -2.52mm) mPFC- and vHC-projecting cells clustered mediodorsally. vHC-projecting cells were located predominantly dorsally while mPFC-projecting cells were more abundant medially. Dual-projecting cells were preferentially situated in midline regions and, in general, were not observed in lateral portions of PVT. Here, there was also a clear separation in axonal processes from these populations, with mPFC-projecting axons dominating the lateral and ventral portions of the nucleus and vHC-projecting axons confined mediodorsally and immediately ventral to the 3V (Fig. 2Aiii). The dense mesh of axons and puncta from mPFC-projecting cells along the lateral aspects of the nucleus suggests that mPFC-projecting cells exert a strong synaptic influence on other PVT cells.

At the most posterior levels of PVT (AP∼ -3.00mm), where the area of PVT is smaller compared to anterior levels, mPFC-projecting cells were more abundant than vHC-projecting cells, which clustered mainly in midline areas (Fig. 2Aiv). Synaptic puncta from vHC-projecting axon terminals were observed throughout, but predominantly in the dorsal band. mPFC-projecting axons and puncta was also observed throughout, however, found in larger numbers ventrally. Dual-projecting cells were mostly localized in the medial regions and within clusters of mPFC- and vHC-projecting cells (Fig. 2Aiv).

We quantified total cells counts in PVT (DAPI expressing), mPFC-projecting cells (magenta), vHC-projecting cells (green), and dual-projecting cells (white). We found that the percent cell density (# of counted cells/DAPI) was highest for mPFC-projecting cells (Mean = 15.27%, SD = 4.72%; or excluding dual projecting cells from the counts Mean = 13.60%, SD = 4.51%) compared to vHC-projecting population (Mean = 6.95% SD = 2.93%; or excluding dual projecting cells from the counts Mean = 5.27%, SD = 2.65%). Dual projecting neurons accounted for about 1.67% (SD = 0.72) of all cells in PVT (Fig. 2C). A one-way repeated measures ANOVA showed that mean percent cell density comparisons of mPFC-projecting, vHC-projecting, and dual-projectin cells was significantly different (F_(2,30)_ = 82.64, p < 0.0005). Post hoc analysis (with a Bonferroni correction) indicated that there were significant differences in mean cell densities between mPFC-projecting and vHC-projecting, mPFC-projecting and dual-projecting; and vHC-projecting and dual-projecting cells (all p’s < 0.0005), suggesting these projections populations are not uniformly represented in PVT.

When PVT was separated by AP level, the percent cell density of mPFC-projecting cells remained consistently larger than the vHC-projecting cells across all levels (Fig. 2D). Of note, cell densities of both mPFC- and vHC-projecting populations was highest in anterior levels and dropped in posterior levels. Interestingly, dual-projecting cell densities remained relatively constant across all levels but was highest at anterior and posterior levels (Fig. 2D). We evaluated cell density differences among cell populations (mPFC-projecting, vHC-projecting and dual-projecting cells) across the four anterior-posterior levels of PVT (anterior, anteromedial, posteromedial and posterior). One-way repeated measures ANOVAs revealed significant differences in cell density across mPFC-projecting, vHC-projecting and dual projecting cell populations at the anterior (F_(2,6)_ = 15.58, p = 0.004), anteromedial (F_(2,6)_ = 33.79, p = 0.001), posteromedial (F_(2,6)_ = 19.34, p = 0.002), and posterior (F_(2,6)_ = 29.13, p = 0.001) levels. Post hoc analysis (Bonferroni correction) revealed significant differences between mPFC-projecting and dual-projecting cell densities at all levels (all p’s < 0.05); vHC-projecting and dual-projecting cell densities at the anteromedial level (p = 0.045) and between mPFC-projecting and vHC-projecting cell densities at the posterior levels (p = 0.050).

### Dual mPFC-vHC projecting PVT cells do not express CR^+^ or CB^+^

We previously described the topographical localization of dual mPFC-HC projecting neurons in RE, and their relationship to CR^+^ and CB^+^ cells (Viena et al. 2021). Given the similarities found between PVT and RE CR^+^ and CB^+^ cells, we also explored the relationship of PVT dual-projecting cells in relation to CR^+^ and CB^+^ expressing neurons. We found that dual-projecting cells in PVT expressed neither CR^+^ or CB^+^ (Fig. 3Ai-Aiv) similar to RE dual-projecting cells. Also similar to RE, dual-projecting cells were nested within clusters of CR^+^ and CB^+^ cells. We then examined the spatial distribution of CR^+^ or CB^+^ cells relative to dual-projecting cells in PVT subset of cells (n = 41 neurons, 2 animals per condition; Fig. 3Ai-Aiv). We found that there was a progressive increase of CR^+^ or CB^+^ cells (Fig. 3Bi-Biv), and this relationship was linearly correlated in both aPVT (CR^+^ R^2^ = 0.793, CB^+^ cells, R^2^ = 0.833) and pPVT (CR^+^ cells R^2^ = 0.897; CB^+^ cells R^2^ = 0.793; all p < 0.0001; Fig. 3Bi-Biv). While this gave us an understanding of the relationship in terms of distance, we wanted to test if these PVT CR^+^ or CB^+^ cells held any sort of radial organization. Thus, we also examined the average cell area density of CR^+^ or CB^+^ cells per total area in every 10μm radius around dual-projecting cells up to a 100μm total radius (# cells per selected radius area; Fig. 3Ci-Civ). We found that within the first annulus (closest to the PVT dual neurons), the percent cell area density was highest in both CR^+^ and CB^+^ cells in anterior and posterior sections. However, there was a sharp and exponential decrease in the average percent cell area density at each subsequent 10um annulus area from the centroid (dual-projecting cells). We then modeled the radial distribution using both linear and one-phase exponential decay. The exponential decay fit the distributions well: CR^+^: aPVT exponential R^2^ = 0.955 (F_(1,7)_ = 150.0, p < 0.0001) vs. linear R^2^= 0.636 (F_(1,8)_ = 13.98, p = 0.006) ; pPVT exponential R^2^ = 0.865 (F_(1,7)_ = 44.79, p = 0.0003) vs. linear R^2^=0.566 (F_(1,8)_ = 10.43, p = 0.012) ; CB^+^ aPVT exponential R^2^ = 0.979 (F_(1,7)_ = 320.9, p < 0.0001) vs. linear 0.570 (F_(1,8)_ = 10.61, p = 0.012); pPVT exponential R^2^ = 0.964 (F_(1,7)_ =187.0, p < 0.0001) vs. linear R^2^= 0.469 (F_(1,8)_ = 7.069, p = 0.029) . We confirmed that the exponential decay model fit significantly better through direct comparison: CR^+^ aPVT (F_(1,7)_ = 50.13, p = 0.0002), CR^+^ pPVT (F_(1,7)_ = 15.48, p = 0.0056) ; CB^+^ aPVT (F_(1,7)_ = 134.0, p < 0.0001), CB^+^ pPVT (F_(1,7)_ = 96.00, p < 0.0001) ; Figs. Ci-Civ). Overall, these results suggest clusters of CR^+^ or CB^+^ around dual mPFC-vHC projecting cells in PVT that may serve as functional zones.

**Figure 3.**
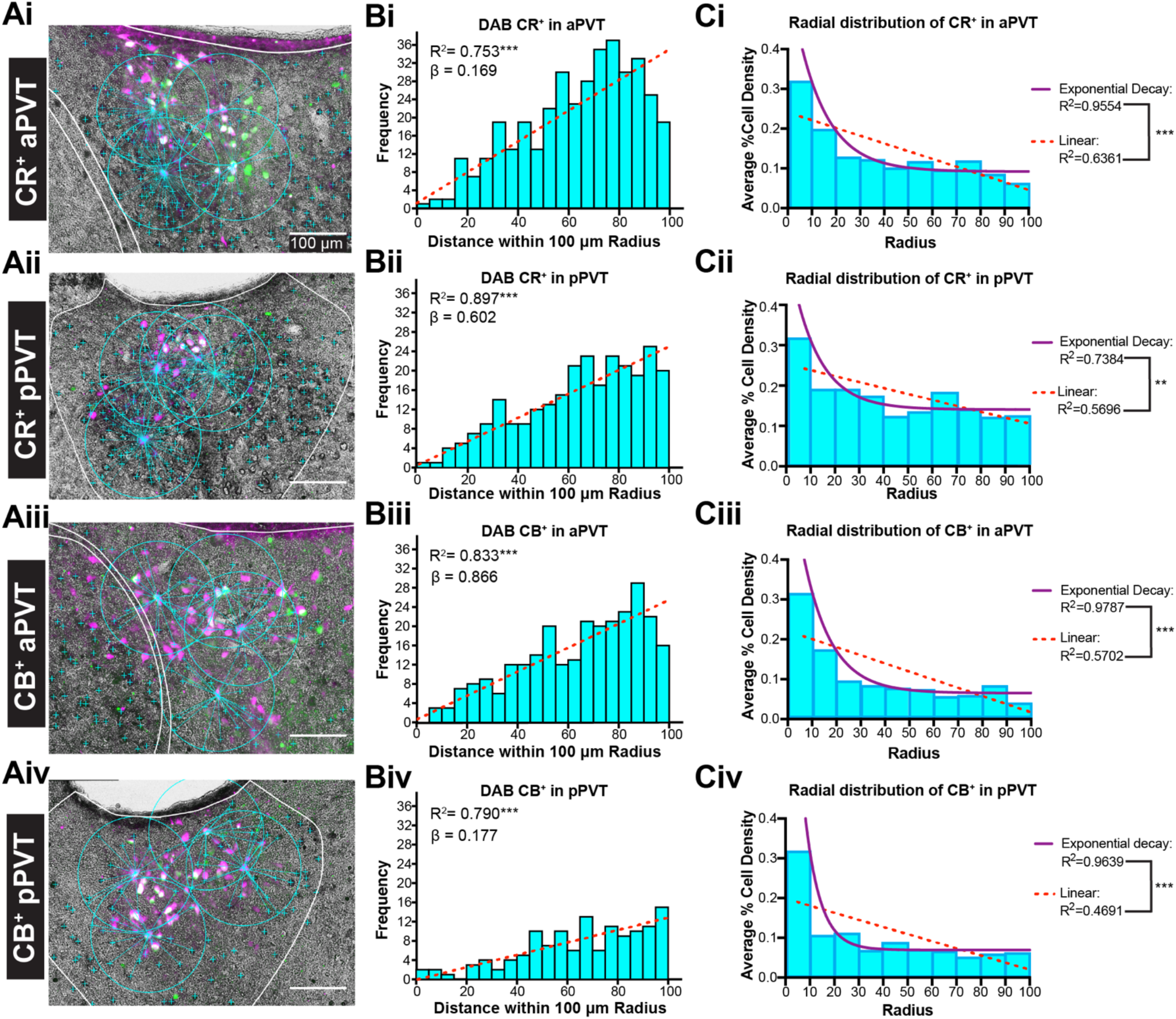
Calretinin and calbindin are not expressed in dual mPFC-vHC projecting cells in PVT. A) Merged captures of immunofluorescent PVT retrogradely labeled mPFC-vHC (dual) cells and DAB CR^+^ (Ai-Aii) or DAB CB^+^ (Aiii-Aiv) cells in anterior and posterior sections. The relative distance between the centroid of the dual labeled cell in respect to DAB CR^+^ or CB^+^ stained cell bodies (cyan “+” signs), measured within 100 um radiuses. No co-expression of either calcium binding protein was found on dual projecting cells. B) Frequency distribution of PVT DAB CR^+^ or CB^+^ cell counts in rostral and caudal PVT relative to distance from radius (5 um bins) from PVT dual (white) labeled cells. C) PVT CR+ or CB+ average cell area density (total area every 10 um^2^ radius within a 100um radius from the center of dual mPFC-HC projecting PVT cells). Linear and exponential decay lines were fitted to model the random distribution of CR^+^ and CB^+^ clusters. Abbreviations: aPVT, anterior PVT; CB, calretinin; CR, calbindin; DAB, 3,3′-Diaminobenzidine; HC, hippocampus mPFC; medial prefrontal cortex; PVT, paraventricular nucleus of the thalamus; pPVT, posterior PVT; vHC, ventral hippocampus.

## Discussion

### Three distinct PVT-projecting cell populations: mPFC-projecting, vHC-projecting and dual mPFC-vHC projecting

The present study provides a description of three distinct types of PVT neuronal populations, mPFC-projecting, vHC-projecting and dual mPFC-vHC projecting cells, which were quantified across its anteroposterior axis (see Fig. 1E for a summary diagram). In line with previous studies, we found a large proportion (∼15%) of PVT neurons that projected to prelimbic and infralimbic regions (layers V/VI) of the rodent mPFC but not vHC (Li and Kirouac 2008; Vertes and Hoover 2008). Additionally, we described a population of PVT neurons that project to ventral CA1 (∼7%), but not to PL/IL. Lastly, we describe a new projection population class of PVT cells that send collaterals to both mPFC and vHC (∼2%). Previous reports have described projections from PVT to ventral CA1 (Su and Bentivoglio 1990) and ventral subiculum (Moga et al. 1995; Li and Kirouac 2008; Vertes and Hoover 2008), but this is the first study to identify dual mPFC-vHC projecting cells in PVT. Similar dual-projecting neurons have been described in RE of the midline thalamus (Hoover and Vertes 2012; Varela et al. 2014; Viena et al. 2021). We also found topographically arranged axonal distribution of these neurons throughout PVT such that mPFC projections tend to be dorsolateral and HC projections tend to be dorsomedial. These findings suggest that PVT, just like RE, influences mPFC-HC interactions at a fine temporal scale of synchrony made possible by dual-projecting neurons, and thus an equally important role in facilitating mPFC-HC interactions.

### PVT dual mPFC-vHC projecting cells display the same characteristics of RE dual mPFC-vHC projecting cells

In our previous report (Viena et al., 2021), we provided a detailed description of PVT and RE neurons expressing calcium biding proteins CR and CB and their relationship to RE dual mPFC-vHC projecting cells. CR^+^ and CB^+^ cells in both RE and PVT shared several characteristics such as location, distribution, and cell size. The noted symmetrical similarities in terms of calcium binding protein expression, have led us to further consider unifying characteristics of these typically segregated midline thalamic nuclei. When we assessed the co-expression between PVT CR^+^ and CB^+^ neurons and PVT dual mPFC-vHC projecting cells, we found that just like in RE, PVT dual-projecting cells did not co-express CR or CB. Similarly, they were surrounded by groups of CR^+^ and CB^+^ cells in the same way. That is, CR^+^ or CB^+^ cells surrounding PVT dual-projecting cells are spatially segregated in clusters that follow a Poisson-like distribution within a 100-µm radius. Generally, this type of radial organization has been described in other thalamic regions (Guillery and Sherman 2002; Sherman 2004; Bickford 2015), but the distribution is strikingly similar in PVT and RE. The organization between CR^+^, CB^+^, and PVT dual-projecting cells may integrate activity from mPFC-dominated subcircuits with vHC-dominated subcircuits thereby triggering dual mPFC-HC projecting cells. Once activated, dual mPFC-HC projecting cells could then send volleys of synchronizing action potentials to both mPFC and HC promoting mPFC-HC interactions under precise physiological activity states.

### Are the dorsal and ventral midline thalamic nuclei best thought of as a single anatomical unit?

The present study adds evidence to the idea that the midline thalamus is best conceived as a single anatomical structure that wraps around the third ventricle with dorsal and ventral portions. We previously based this notion on the distribution of CR+, CB+ and dual CR/CB+ cells (Viena et al., 2021). This unified midline thalamus conceptualization is similar to the hippocampus with dorsal and ventral subregions that differ less by hard anatomical boundaries and architectural features, but more by functional zones defined by differences in the connectivity profiles. Here, we provide more data showing strong similarities suggestive of a unified midline thalamus: First, there is the presence of dual mPFC-HC projecting neurons in PVT. Like RE, these dual projecting neurons are primarily found between the CR and CB rich zones. Second, the PVT dual-projecting cells share the characteristics of RE dual mPFC-HC projecting neurons because they lack CR and CB labelling themselves but are surrounded by both mPFC- and HC-projecting cells. Third, we also observed a similar projection topography in that axons bound for mPFC tend to distribute dorsolaterally, and axons bound for vHC tend to distribute dorsomedially in PVT. Relative to the location of the third ventricle this pattern looks very much like peri-reuniens (laterally) and medial RE that tend to project to mPFC and vHC, respectively. Related to this idea, most studies have explored PVT and RE separately, but some reports have argued these nuclei fit more the description of a unitary and functional “limbic thalamus” (Pereira de Vasconcelos and Cassel 2015; Vertes et al. 2015). Earlier distinctions separated the midline thalamus into four discrete nuclei anatomically based off projection target patterns (Van der Werf, 2002). The unified midline thalamus conceptualization coalesces both of these views.

### Functional significance of PVT dual mPFC-vHC projection populations

PVT cells that send collaterals to both mPFC and vHC are among many other collateralized cell types in PVT with multiple downstream targets such as PL-NAc (Bubser and Deutch 1998; Otake and Nakamura 1998), PL-amygdala (Dong et al. 2017), BLA-NAc (Dong et al. 2017). While the relationship between dual mPFC-HC projecting PVT cells and these other projection targets is not known, it does speak to PVT as a unique node in integrating areas processing executive functions, motivated behaviors, salient cues, and memory (Dong et al. 2017; Zhu et al. 2018). For example, the role of PVT in the PL-CEa circuit has been explored as a key player in long-term memory expression critical for fear retrieval (Padilla-Coreano et al. 2012; Do-Monte et al. 2015; Penzo et al. 2015). Additionally, the radial distribution of CR^+^ and CB^+^ with respect to dual projecting-cells suggests that there are well-organized zones that potentially integrate information from subsets of the many brain regions that innervate PVT. With dual projecting neurons throughout, this can then cause highly synchronous changes to mPFC-HC activity states/modes. However, this does not address many of the better studied PVT network relationships. It would be beneficial to study PVT further in the context of cortico-limbic networks separating mPFC-projecting, vHC-projecting and dual mPFC-vHC projecting neurons in PVT. For example, how does each population influence activity in the mPFC→CEa or mPFC→NAc circuitry. Further, investigating how PVT is involved in the modulation of activity within these circuits (e.g., looking at coherent delta and theta rhythms; Schultheiss et al., 2020) within the mPFC-HC system may help elucidate PVT’s role in drug addiction and relapse, for example using cell-type specific activation approaches.

### PVT→PVT projections

The present findings also suggest a topographically arranged intra-PVT projection patterns based on the dense collaterals and the mesh of florescent puncta (also see Vertes & Hoover, 2008). Although more studies are needed to characterize these intra-PVT connections, their presence suggests that PVT neurons projecting to mPFC and vHC can also influence local PVT network activity through PVT→PVT interactions. Speculatively, these interactions would provide feedback excitation to potentially boost activity to mPFC-HC mode setting, refine weakly dominant representations to more dominant ones, or potentially trigger different cascades of network states (e.g., mPFC-HC directionality by rocking back and forth between projection populations). Input-output tracing investigations (Gergues et al. 2020) has provided some details about how PVT differentially innervates multiple vCA1 circuits including vCA1→mPFC, vCA1→basolateral amygdala, and vCA1→lateral habenula. Settling on a dominant circuit for any given behavior in this network may be boosted by this local feedback, but testing this hypothesis will require future studies using sophisticated projection-specific cell methods in combination with optogenetics and electrophysiological measures.

## Conclusion

We provide a comprehensive anatomical description of three distinct PVT cell populations potentially associated to the mPFC-HC circuity: mPFC-projecting, vHC-projecting, and mPFC-vHC projecting. Further studies will be needed to identifying the functional roles that these different PVT cell populations play the mPFC-HC network. Ultimately, this will help elucidate the mechanisms regulated by PVT that go awry in mental health disorders such as post-traumatic stress disorders and drug addiction.

## Acknowledgements

The authors would like to thank Daniela Silva, Nashya Linares, and Amanda Pacheco Spiezwak for their assistance in cell counting. This work was supported in part by NIH grant R01 MH113626S1 to T.A.A., and funds from the FIU CASE Distinguished Postdoctoral Program to T.D.V.

## Author Contributions

Experimental design: T.D.V., T.A.A. Performed research, drafted manuscript and analyzed results: T.D.V., G.E.R. Edited the manuscript: T.D.V., G.E.R. and T.A.A.

## Declaration of Interests

The authors declare no competing interests.

## Notes

### Competing Interest Statement

The authors have declared no competing interest.

## References

Aggleton JP, Brown MW (1999) Episodic memory, amnesia, and the hippocampal–anterior thalamic axis. Behavioral and Brain Sciences 22:425–444. https://doi.org/10.1017/S0140525X99002034

Avigan PD, Cammack K, Shapiro ML (2020) Flexible spatial learning requires both the dorsal and ventral hippocampus and their functional interactions with the prefrontal cortex. Hippocampus 30:733–744. https://doi.org/10.1002/hipo.23198

Bannerman DM, Grubb M, Deacon RMJ, et al (2003) Ventral hippocampal lesions affect anxiety but not spatial learning. Behav Brain Res 139:197–213. https://doi.org/10.1016/s0166-4328(02)00268-1

Bickford ME (2015) Thalamic Circuit Diversity: Modulation of the Driver/Modulator Framework. Front Neural Circuits 9:86. https://doi.org/10.3389/fncir.2015.00086

Bubser M, Deutch AY (1998) Thalamic paraventricular nucleus neurons collateralize to innervate the prefrontal cortex and nucleus accumbens. Brain Res 787:304–310. https://doi.org/10.1016/s0006-8993(97)01373-5

Cassel J-C, Ferraris M, Quilichini P, et al (2021) The reuniens and rhomboid nuclei of the thalamus: A crossroads for cognition-relevant information processing? Neurosci Biobehav Rev 126:338–360. https://doi.org/10.1016/j.neubiorev.2021.03.023

Cassel J-C, Pereira de Vasconcelos A, Loureiro M, et al (2013) The reuniens and rhomboid nuclei: neuroanatomy, electrophysiological characteristics and behavioral implications. Prog Neurobiol 111:34–52. https://doi.org/10.1016/j.pneurobio.2013.08.006

Choi EA, Jean-Richard-dit-Bressel P, Clifford CWG, McNally GP (2019) Paraventricular Thalamus Controls Behavior during Motivational Conflict. J Neurosci 39:4945–4958. https://doi.org/10.1523/JNEUROSCI.2480-18.2019

Choi EA, McNally GP (2017) Paraventricular Thalamus Balances Danger and Reward. J Neurosci 37:3018–3029. https://doi.org/10.1523/JNEUROSCI.3320-16.2017

Cholvin T, Loureiro M, Cassel R, et al (2013) The Ventral Midline Thalamus Contributes to Strategy Shifting in a Memory Task Requiring Both Prefrontal Cortical and Hippocampal Functions. J Neurosci 33:8772–8783. https://doi.org/10.1523/JNEUROSCI.0771-13.2013

Churchwell JC, Kesner RP (2011) Hippocampal-prefrontal dynamics in spatial working memory: Interactions and independent parallel processing. Behav Brain Res 225:389–395. https://doi.org/10.1016/j.bbr.2011.07.045

Dolleman-van der Weel MJ, Griffin AL, Ito HT, et al (2019) The nucleus reuniens of the thalamus sits at the nexus of a hippocampus and medial prefrontal cortex circuit enabling memory and behavior. Learn Mem 26:191–205. https://doi.org/10.1101/lm.048389.118

Do-Monte FH, Quiñones-Laracuente K, Quirk GJ (2015) A temporal shift in the circuits mediating retrieval of fear memory. Nature 519:460–463. https://doi.org/10.1038/nature14030

Dong X, Li S, Kirouac GJ (2017) Collateralization of projections from the paraventricular nucleus of the thalamus to the nucleus accumbens, bed nucleus of the stria terminalis, and central nucleus of the amygdala. Brain Struct Funct 222:3927–3943. https://doi.org/10.1007/s00429-017-1445-8

Dong X, Li Y, Kirouac GJ (2015) Blocking of orexin receptors in the paraventricular nucleus of the thalamus has no effect on the expression of conditioned fear in rats. Front Behav Neurosci 9:. https://doi.org/10.3389/fnbeh.2015.00161

Eichenbaum H (2017) Prefrontal-hippocampal interactions in episodic memory. Nat Rev Neurosci 18:547–558. https://doi.org/10.1038/nrn.2017.74

Ferraris M, Ghestem A, Vicente AF, et al (2018) The Nucleus Reuniens Controls Long-Range Hippocampo– Prefrontal Gamma Synchronization during Slow Oscillations. J Neurosci 38:3026–3038. https://doi.org/10.1523/JNEUROSCI.3058-17.2018

Gao C, Leng Y, Ma J, et al (2020) Two genetically, anatomically and functionally distinct cell types segregate across anteroposterior axis of paraventricular thalamus. Nature Neuroscience 23:217–228. https://doi.org/10.1038/s41593-019-0572-3

Gergues MM, Han KJ, Choi HS, et al (2020) Circuit and molecular architecture of a ventral hippocampal network. Nature Neuroscience 23:1444–1452. https://doi.org/10.1038/s41593-020-0705-8

Groenewegen HJ, Berendse HW (1994) The specificity of the ‘nonspecific’ midline and intralaminar thalamic nuclei. Trends in Neurosciences 17:52–57. https://doi.org/10.1016/0166-2236(94)90074-4

Guillery RW, Sherman SM (2002) Thalamic relay functions and their role in corticocortical communication: generalizations from the visual system. Neuron 33:163–175. https://doi.org/10.1016/s0896-6273(01)00582-7

Haight JL, Flagel SB (2014) A potential role for the paraventricular nucleus of the thalamus in mediating individual variation in Pavlovian conditioned responses. Front Behav Neurosci 8:. https://doi.org/10.3389/fnbeh.2014.00079

Hallock HL, Wang A, Griffin AL (2016) Ventral Midline Thalamus Is Critical for Hippocampal–Prefrontal Synchrony and Spatial Working Memory. J Neurosci 36:8372–8389. https://doi.org/10.1523/JNEUROSCI.0991-16.2016

Hauer BE, Pagliardini S, Dickson CT (2019) The Reuniens Nucleus of the Thalamus Has an Essential Role in Coordinating Slow-Wave Activity between Neocortex and Hippocampus. eNeuro 6:ENEURO.0365-19.2019. https://doi.org/10.1523/ENEURO.0365-19.2019

Heydendael W, Sharma K, Iyer V, et al (2011) Orexins/Hypocretins Act in the Posterior Paraventricular Thalamic Nucleus During Repeated Stress to Regulate Facilitation to Novel Stress. Endocrinology 152:4738–4752. https://doi.org/10.1210/en.2011-1652

Hill-Bowen LD, Flannery JS, Poudel R (2020) Paraventricular Thalamus Activity during Motivational Conflict Highlights the Nucleus as a Potential Constituent in the Neurocircuitry of Addiction. J Neurosci 40:726–728. https://doi.org/10.1523/JNEUROSCI.1945-19.2019

Hoover WB, Vertes RP (2012) Collateral projections from nucleus reuniens of thalamus to hippocampus and medial prefrontal cortex in the rat: a single and double retrograde fluorescent labeling study. Brain Struct Funct 217:191–209. https://doi.org/10.1007/s00429-011-0345-6

Huang H, Ghosh P, van den Pol AN (2006) Prefrontal Cortex–Projecting Glutamatergic Thalamic Paraventricular Nucleus-Excited by Hypocretin: A Feedforward Circuit That May Enhance Cognitive Arousal. Journal of Neurophysiology 95:1656–1668. https://doi.org/10.1152/jn.00927.2005

Ito HT, Zhang S-J, Witter MP, et al (2015) A prefrontal–thalamo–hippocampal circuit for goal-directed spatial navigation. Nature 522:50–55. https://doi.org/10.1038/nature14396

James MH, Charnley JL, Jones E, et al (2010) Cocaine- and Amphetamine-Regulated Transcript (CART) Signaling within the Paraventricular Thalamus Modulates Cocaine-Seeking Behaviour. PLOS ONE 5:e12980. https://doi.org/10.1371/journal.pone.0012980

Jayachandran M, Linley SB, Schlecht M, et al (2019) Prefrontal Pathways Provide Top-Down Control of Memory for Sequences of Events. Cell Reports 28:640-654.e6. https://doi.org/10.1016/j.celrep.2019.06.053

Jones MW, Wilson MA (2005) Theta Rhythms Coordinate Hippocampal–Prefrontal Interactions in a Spatial Memory Task. PLoS Biol 3:. https://doi.org/10.1371/journal.pbio.0030402

Kirouac GJ (2015) Placing the paraventricular nucleus of the thalamus within the brain circuits that control behavior. Neuroscience & Biobehavioral Reviews 56:315–329. https://doi.org/10.1016/j.neubiorev.2015.08.005

Li S, Kirouac GJ (2012) Sources of inputs to the anterior and posterior aspects of the paraventricular nucleus of the thalamus. Brain Struct Funct 217:257–273. https://doi.org/10.1007/s00429-011-0360-7

Li S, Kirouac GJ (2008) Projections from the paraventricular nucleus of the thalamus to the forebrain, with special emphasis on the extended amygdala. J Comp Neurol 506:263–287. https://doi.org/10.1002/cne.21502

Martin-Fardon R, Boutrel B (2012) Orexin/hypocretin (Orx/Hcrt) transmission and drug-seeking behavior: is the paraventricular nucleus of the thalamus (PVT) part of the drug seeking circuitry? Frontiers in Behavioral Neuroscience 6:75. https://doi.org/10.3389/fnbeh.2012.00075

Moga MM, Weis RP, Moore RY (1995) Efferent projections of the paraventricular thalamic nucleus in the rat. Journal of Comparative Neurology 359:221–238. https://doi.org/10.1002/cne.903590204

Moser M-B, Moser EI (1998) Functional differentiation in the hippocampus. Hippocampus 8:608–619. https://doi.org/10.1002/(SICI)1098-1063(1998)8:6<608::AID-HIPO3>3.0.CO;2-7

Neumann PA, Wang Y, Yan Y, et al (2016) Cocaine-Induced Synaptic Alterations in Thalamus to Nucleus Accumbens Projection. Neuropsychopharmacology 41:2399–2410. https://doi.org/10.1038/npp.2016.52

Otake K, Nakamura Y (1998) Single midline thalamic neurons projecting to both the ventral striatum and the prefrontal cortex in the rat. Neuroscience 86:635–649. https://doi.org/10.1016/s0306-4522(98)00062-1

Padilla-Coreano N, Do-Monte FH, Quirk GJ (2012) A time-dependent role of midline thalamic nuclei in the retrieval of fear memory. Neuropharmacology 62:457–463. https://doi.org/10.1016/j.neuropharm.2011.08.037

Penzo MA, Robert V, Tucciarone J, et al (2015) The paraventricular thalamus controls a central amygdala fear circuit. Nature 519:455–459. https://doi.org/10.1038/nature13978

Pereira de Vasconcelos A, Cassel J-C (2015) The nonspecific thalamus: A place in a wedding bed for making memories last? Neuroscience & Biobehavioral Reviews 54:175–196. https://doi.org/10.1016/j.neubiorev.2014.10.021

Schultheiss NW, Schlecht M, Jayachandran M, et al (2020) Awake delta and theta-rhythmic hippocampal network modes during intermittent locomotor behaviors in the rat. Behavioral Neuroscience 134:529–546. https://doi.org/10.1037/bne0000409

Sherman SM (2004) Interneurons and triadic circuitry of the thalamus. Trends Neurosci 27:670–675. https://doi.org/10.1016/j.tins.2004.08.003

Squire LR, Stark CEL, Clark RE (2004) The Medial Temporal Lobe. Annual Review of Neuroscience 27:279–306. https://doi.org/10.1146/annurev.neuro.27.070203.144130

Su HS, Bentivoglio M (1990) Thalamic midline cell populations projecting to the nucleus accumbens, amygdala, and hippocampus in the rat. J Comp Neurol 297:582–593. https://doi.org/10.1002/cne.902970410

Van der Werf YD, Witter MP, Groenewegen HJ (2002) The intralaminar and midline nuclei of the thalamus. Anatomical and functional evidence for participation in processes of arousal and awareness. Brain Research Reviews 39:107–140. https://doi.org/10.1016/S0165-0173(02)00181-9

Varela C, Kumar S, Yang JY, Wilson MA (2014) Anatomical substrates for direct interactions between hippocampus, medial prefrontal cortex, and the thalamic nucleus reuniens. Brain Struct Funct 219:911–929. https://doi.org/10.1007/s00429-013-0543-5

Vertes RP, Hoover WB (2008) Projections of the paraventricular and paratenial nuclei of the dorsal midline thalamus in the rat. J Comp Neurol 508:212–237. https://doi.org/10.1002/cne.21679

Vertes RP, Hoover WB, Szigeti-Buck K, Leranth C (2007) Nucleus reuniens of the midline thalamus: Link between the medial prefrontal cortex and the hippocampus. Brain Research Bulletin 71:601–609. https://doi.org/10.1016/j.brainresbull.2006.12.002

Vertes RP, Linley SB, Hoover WB (2015) Limbic circuitry of the midline thalamus. Neuroscience & Biobehavioral Reviews 54:89–107. https://doi.org/10.1016/j.neubiorev.2015.01.014

Viena TD, Linley SB, Vertes RP (2018) Inactivation of nucleus reuniens impairs spatial working memory and behavioral flexibility in the rat. Hippocampus 28:297–311. https://doi.org/10.1002/hipo.22831

Viena TD, Rasch GE, Silva D, Allen TA (2021) Calretinin and calbindin architecture of the midline thalamus associated with prefrontal–hippocampal circuitry. Hippocampus 31:770–789. https://doi.org/10.1002/hipo.23271

Wirt RA, Hyman JM (2017) Integrating Spatial Working Memory and Remote Memory: Interactions between the Medial Prefrontal Cortex and Hippocampus. Brain Sci 7:. https://doi.org/10.3390/brainsci7040043

Xu C, Krabbe S, Gründemann J, et al (2016) Distinct Hippocampal Pathways Mediate Dissociable Roles of Context in Memory Retrieval. Cell 167:961-972.e16. https://doi.org/10.1016/j.cell.2016.09.051

Yoon T, Okada J, Jung MW, Kim JJ (2008) Prefrontal cortex and hippocampus subserve different components of working memory in rats. Learn Mem 15:97–105. https://doi.org/10.1101/lm.850808

Zhu Y, Nachtrab G, Keyes PC, et al (2018) Dynamic Salience Processing in Paraventricular Thalamus Gates Associative Learning. Science 362:423–429. https://doi.org/10.1126/science.aat0481

Zhu Y, Wienecke CFR, Nachtrab G, Chen X (2016) A thalamic input to the nucleus accumbens mediates opiate dependence. Nature 530:219–222. https://doi.org/10.1038/nature16954

Zimmerman EC, Grace AA (2016) The Nucleus Reuniens of the Midline Thalamus Gates Prefrontal-Hippocampal Modulation of Ventral Tegmental Area Dopamine Neuron Activity. J Neurosci 36:8977–8984. https://doi.org/10.1523/JNEUROSCI.1402-16.2016

